# Rab40 GTPases regulate AMBRA1-mediated transcription and cell migration

**DOI:** 10.1101/2024.11.07.622540

**Authors:** Revathi Sampath, Katherine Vaeth, Valeryia Mikalayeva, Vytenis Arvydas Skeberdis, Rytis Prekeris, Ke-Jun Han

## Abstract

The Rab40 subfamily are unique small monomeric GTPases that form CRL5-based ubiquitin E3 ligase complex and regulate ubiquitylation of specific target proteins. Recent studies have shown that Rab40s play an important role in regulating cell migration, but the underlying mechanisms of Rab40/CRL5 complex function are still not fully understood. In this study we identified AMBRA1 as a novel binding partner of Rab40 GTPases and showed that this interaction mediates a bi-directional crosstalk between CRL4 and CRL5 E3 ligases. Importantly, we found that Rab40/CRL5 ubiquitylates AMBRA1, which does not result in AMBRA1 degradation, but instead it seems to induce AMBRA1-dependent regulation of gene transcription. The global transcriptional profiles identified by RNA-seq showed that AMBRA1 regulates transcription of genes related to cell adhesion and migration. Additionally, we have shown that AMBRA1-dependent transcription regulation does not require the enzymatic activity of AMBRA1/CRL4, and that Rab40-induced AMBRA1 ubiquitylation leads to dissociation of AMBRA1/CRL4 complex. Taken together, our findings reveal a novel function of Rab40/CRL5 complex as an important regulator for AMBRA1-dependent transcription of genes involved in cell migration.

## INTRODUCTION

Rab proteins are small monomeric GTPases belonging to the Ras GTPase superfamily. Rab GTPases are evolutionarily conserved and function as key regulators of eukaryotic membrane trafficking. The human genome includes over 70 Rab GTPases, which can be divided into 10 major subfamilies [1–4]. Among them, Rab40 subfamily is unique because it has an extended C-terminal, which contains a suppressor of cytokine signaling (SOCS) box motif [5, 6], thus mediates interaction with Cullin5 to form an E3 ubiquitin ligase complex (Rab40/CRL5) [5–7]. Therefore, Rab40s function not only as molecular regulators of membrane traffic, but also as ubiquitin E3 ligase complex to mediate target protein ubiquitylation [5–7].

The Rab40 subfamily consists of four closely related proteins: Rab40a, Rab40al, Rab40b, and Rab40c [4, 5, 7]. We and others previously demonstrated that Rab40a and Rab40b are required for regulating cancer cell migration and invasion by promoting extracellular matrix degradation, dynamics of focal adhesion sites (FAs), and invadopodia formation [8–13]. Specifically, Rab40a was reported to mediate proteasomal degradation of RhoU whilst Rab40b ubiquitylate Eplin and Rap2, thus promoting cell migration by altering FA dynamics and stress fiber formation [8, 10, 12]. Additionally, we have shown that Rab40c binds the protein phosphatase 6 (PP6) complex and ubiquitylates ANKRD28 subunit, thus leading to its lysosomal degradation which ultimately also affects FAs [11]. All these findings suggest that Rab40 subfamily GTPases may have evolved to regulate actin dynamics and FA turnover by mediating ubiquitylation of a specific subset of proteins; however, it remains to be fully understood how Rab40 function is regulated and what molecular machinery governs Rab40-dependent cell migration and invasion.

Our recent proteomics screen identified AMBRA1 (activating molecule in Beclin1-regulated autophagy) as a putative target for Rab40c/CRL5-dependent ubiquitylation [11]. AMBRA1 is a WD40 domain-containing protein and is involved in various biological processes including autophagy and cell division [14–18]. AMBRA1 acts as a substrate-recognition component of a Cullin4-DDB1 E3 ubiquitin-protein ligase complex (AMBRA1/CRL4), promoting the ubiquitination of Beclin1 and ULK1, and therefore is a key regulator of autophagy [14, 16, 17, 19–21]. Interestingly, it was suggested that AMBRA1 can also mediate crosstalk between Cullin4 and Cullin5-depedent E3 ubiquitin ligases (CRL4 and CRL5) [22, 23]. Under normal conditions, AMBRA1binds to CRL4 and is targeted for proteasomal degradation, presumably a consequence of CRL4-dependent auto-ubiquitylation. It was suggested that upon activation of the autophagy, AMBRA1 inhibits CRL5 activity either by disruption of Elongin-B/Cullin5 (part of CRL5 complex) interaction or by mediating proteasomal degradation of Elongin-C [22, 23]. However, it currently remains unclear how this CRL4 and CRL5 crosstalk is regulated and whether AMBRA1 may have other CRL4-independent functions.

In addition to its role in autophagy, recent research highlights the important role of AMBRA1 in cancer cell migration and proliferation. AMBRA1 is proposed to be a tumor suppressor, and loss of AMBRA1 promotes cancer cell growth and invasion [16, 24–26]. The AMBRA1/CRL4 complex binds to cyclin D, leading to cyclin D ubiquitination and subsequent proteasomal degradation, thereby controlling G1-to-S transition and cell division [15, 27–30]. AMBRA1 also regulates Src activity and Src/focal adhesion kinase (FAK)-mediated cancer cell invasion and migration [31, 32]. AMBRA1 can be recruited to FAs, where it was suggested to bind to both FAK and Src and AMBRA1 removes active phospho-Src from FAs and transports it into autophagic structures, likely for degradation [31, 32].

Since our recent work suggested that AMBRA1 may interact with Rab40c, in this study we decided to investigate whether the interaction between Rab40c and AMBRA1 is a new potential regulatory crosstalk pathway between CRL4 with CRL5 complexes. Consistent with this hypothesis, we show that AMBRA1 enhances Rab40c binding to Cullin5, and that depletion of AMBRA1 increases Rab40c mRNA and protein levels. We also found that loss of AMBRA1 in MDA-MB-231 cells alters FA distribution and promotes cell migration. Intriguingly, at least some of the AMBRA1 effects on cell adhesion and migration appear to be mediated by AMBRA1-dependent regulation of expression of a subset of genes. Using SNAI2 as an example, we showed that AMBRA1-dependent transcription regulation is independent of AMBRA1 binding to CRL4, but it appears to be regulated by Rab40/CRL5-induced ubiquitylation. Thus, we uncovered a new CRL4-independent function of AMBRA1 in regulating cell migration by regulating gene expression.

## RESULTS

### Rab40c is an AMBRA1-binding protein

All Rab40 subfamily of proteins (Rab40a, Rab40b, and Rab40c) regulate cell migration by forming Cullin5-containing Rab40/CRL5 complex and mediating ubiquitylation of the specific target proteins [5–7, 10]. Recent studies from our and other laboratories have demonstrated that Rab40a and Rab40b are both required for cell migration and function by ubiquitylating Eplin, Rap2, and RhoU [8, 10, 12]. In contrast, how Rab40c regulates cell migration remains largely unclear. Thus, in this study we set out to identify new Rab40c/CRL5 substrates. To this end, we developed a proteomics-based screen that relies on our previous studies showing that mutating of the 211-LPLP-216 (to 211-AAAA-216) domain within SOCS box of Rab40 subfamily members (FLAG-Rab40-4A) lead to the decrease in Cullin5 binding and an increase in Rab40 association with its ubiquitylation substrates [11]. Among these putative Rab40c/CRL5 substrate proteins, AMBRA1 was present in the FLAG-Rab40c-4A but not in the FLAG-Rab40c immunoprecipitates (Fig. 1A) [11]. To confirm that AMBRA1 binds to Rab40c, we immunoprecipitated endogenous Rab40c from MDA-MB-231 cells and found that endogenous AMBRA1 co-immunoprecipitated with Rab40c (Fig. 1B). To further confirm Rab40c and AMBRA1 interaction we overexpressed either FLAG-Rab40c or FLAG-Rab40c-4A in 293T cells, followed by precipitation with anti-FLAG antibodies and immunoblotting for endogenous AMBRA1. Consistent with our proteomics data, we found that AMBRA1 predominantly binds to FLAG-Rab40c-4A, although some AMBRA1 could also be detected in wild-type FLAG-Rab40c immunoprecipitate (Fig. 1C). We next tested whether AMBRA1 can also interact with other Rab40 GTPase subfamily members. As shown in the figure 1D, FLAG-tagged Rab40a, Rab40al, and Rab40b all co-immunoprecipitated with endogenous AMBRA1, suggesting AMBRA1 can interact with all the members of the Rab40 subfamily.

**Figure 1.**
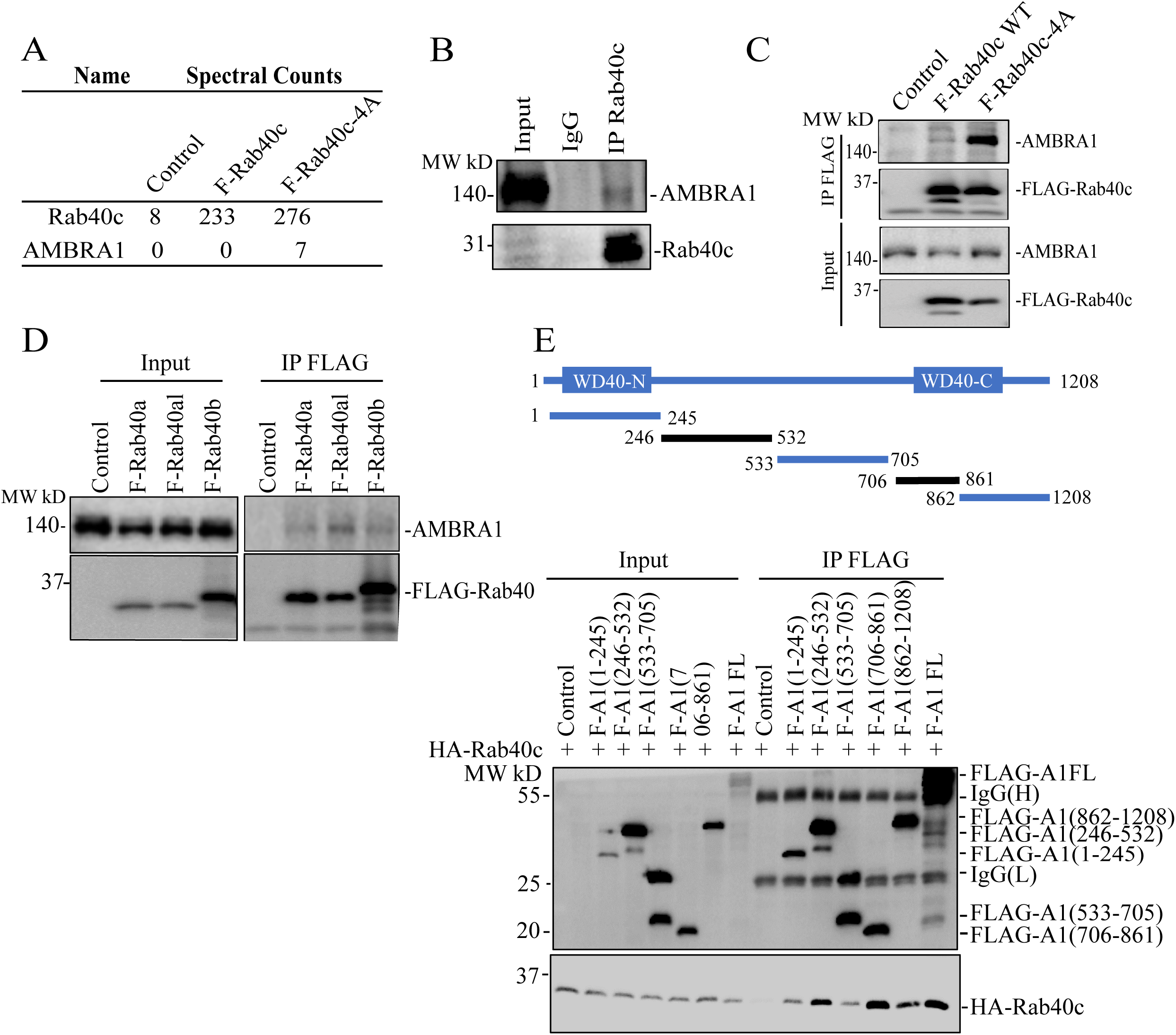
Rab40 subfamily of small monomeric GTPases interact with AMBRA1. (A) AMBRA1 co-immunoprecipiates with FLAG-Rab40c-4A in MDA-MB-231 cells. (B) Rab40c was immunoprecipitated from MDA-MB-231 lysates using anti-Rab40c antibodies. Immunoprecipitate was then blotted with anti-AMBRA1 and anti-Rab40c antibodies. (C) 293T cells were transfected with empty plasmid (control), FLAG-Rab40c, or FLAG-Rab40c-4A plasmids and then FLAG-Rab40c (wild type or 4A mutant) was immunoprecipitated with an anti-FLAG antibody. The precipitates were then blotted with anti-AMBRA1antibodies. (D) 293T cells were transfected with empty vector (control), FLAG-Rab40a, FLAG-Rab40al, or FLAG-Rab40b plasmids and then FLAG-tagged Rab40a/b/c were immunoprecipitated with an anti-FLAG antibody. Cell lysates and precipitates were then blotted with anti-AMBRA1or anti-FLAG antibodies. (E) Top panels: a schematic diagram of AMBRA1 deletion mutants. Lower panels: 293T cells were co-transfected with HA-Rab40c and empty vector or one of FLAG-tagged AMBRA1 deletion mutants. The FLAG-tagged truncation AMBRA1 mutants were then immunoprecipitated with an anti-FLAG antibody and precipitates were blotted with anti-HA antibodies.

We next set out to map which region of AMBRA1 is responsible for Rab40 binding. To this end, we generated a series of FLAG-tagged AMBRA1 deletion mutants including AMBRA1(1-245), (246-532), (533-705), (706-861), and (862-1208) (Fig. 1E), then individually co-transfecting all these constructs with HA-Rab40c into 293T cells, followed by immunoprecipitation with anti-FLAG antibodies. As shown in the figure 1D, FLAG-Rab40c predominately co-precipitated with AMBRA1(246-532) and AMBRA1(706-861), suggesting that AMBRA1 may have two distinct Rab40-binding regions (Fig 1E). Interestingly, it was previously shown that AMBRA1 contains split-WD40 domains (WD40-N and WD40-C) located at N- and C-termini of the proteins (Fig 1E) [33]. These two split-WD40 domains interact to form a fully functional WD40 domain that mediates binding to DDB1 to form the AMBRA1/CRL4 complex. Therefore, due to the close proximity of the AMBRA1(246-532) and AMBRA1(706-861) regions to WD40-N and WD40-C, they also likely form a single Rab40-binding interface.

### AMBRA1 suppresses Rab40c expression but stimulates Rab40/CRL5 complex formation

AMBRA1 is a well-established substrate receptor of Cullin4 ubiquitin ligase complex, which targets protein ubiquitination and subsequent degradation [15, 20, 34–36]. Therefore, we hypothesized that AMBRA1 may also ubiquitylate Rab40c, thus leading to its proteasomal degradation (Fig. 2A). To test this, we first generated AMBRA1 knockout MDA-MB-231 cell lines (KO1 and KO2) using CRISPR/Cas-mediated genome editing, which have been validated by genomic sequencing and Western blotting (Fig. S1B and Fig. 2B-C). As shown in the figure 2B, AMBRA1-KO caused an increase in Rab40c protein levels, whereas it had no effect on the total levels of Cullin5. To further confirm that the increase in Rab40c protein levels is caused by depletion of AMBRA1 we re-introduced AMBRA1, whose expression is driven by a doxycycline (dox) inducible promoter, back into an AMBRA1-KO cell lines. As shown in the figure 2C, Rab40c protein levels gradually decreased and correlated with the levels of dox-induced AMBRA1 expression, thus, suggesting that Rab40c protein level changes are AMBRA1-dependent. Next, we investigated the mechanisms by which AMBRA1 regulates Rab40c protein levels by examining whether AMBRA1 targets Rab40c for proteasomal or lysosomal degradation. To this end, we treated control and AMBA1-KO cells with either the proteasomal inhibitor MG132 or lysosomal inhibitor bafilomycin A1 (BFM). As shown in the figure 2D, MG132 treatment, but not BFM, increased protein levels of Rab40c in both control and AMBRA1-KO cells, suggesting that Rab40c can be degraded by the proteasome pathway. However, the difference in Rab40c protein levels between control and AMBRA1-KO cells did not significantly change after MG132 treatment as compared with the DMSO control. Therefore, the inhibition of proteasomal degradation is insufficient to block AMBRA1-induced increase in Rab40c protein levels. Based on these data, we hypothesized that AMBRA1 may affect transcription of Rab40c mRNA. Consistent with this hypothesis, we found that Rab40c mRNA levels in AMBRA1-KO cells were significantly increased as quantified by real-time quantitative PCR (qRT-PCR) (Fig. 2E).

**Figure 2.**
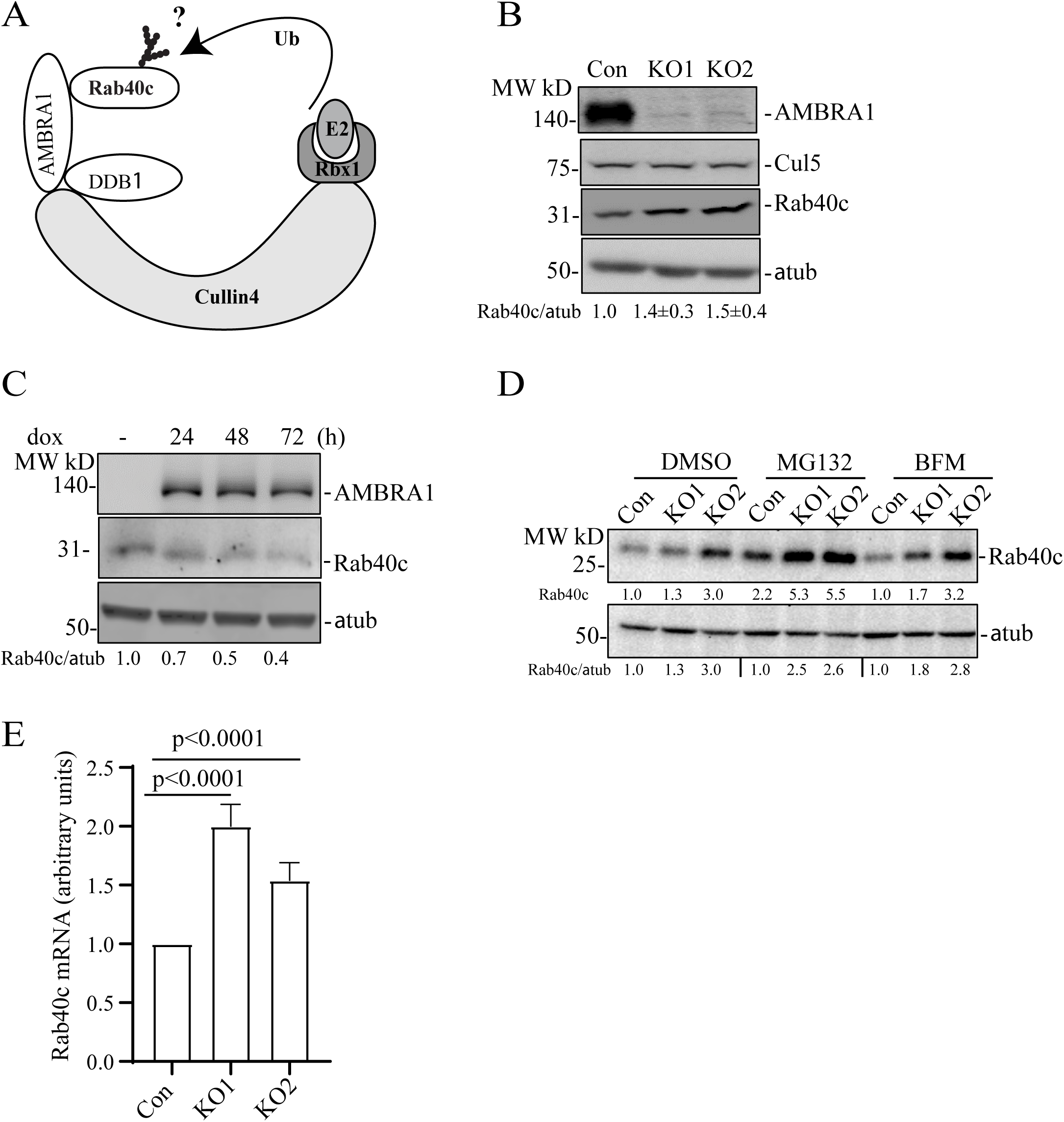
AMBRA1 regulates Rab40c expression. (A) A proposed model showing that Rab40c is a putative substrate of CRL4/AMBRA1 E3 ligase. Ub, ubiquitin. (B) Western blotting analysis of cell lysates from control and AMBRA1-KO cells using indicated antibodies. Relative Rab40c levels in AMBRA1-KO cells were normalized to loading control and control cells. The numbers shown below the blot are the means and SEM derived from three independent experiments. (C) AMBRA1-KO cells expressing AMBRA1 under Tet-On promoter were incubated with 100 ng/ml doxycycline for various time periods and cell lysates were blotted using indicated antibodies. The numbers shown below the blot are relative amounts of Rab40c normalized to the loading control α-tubulin. (D) Control and AMBRA1-KO cells were treated with DMSO, MG132, and Bafilomycin A1. The whole-cell lysates were then analyzed by western blot using indicated antibodies. The numbers shown below the blot are relative amounts of Rab40c normalized to the control samples. (E) qRT-PCR analysis of Rab40c mRNA levels in control and AMBRA1-KO MDA-MB-231 cells. The data shown are the means and SEM derived from three independent biological replicates.

Since AMBRA1 binds to Cullin4 and acts as substrate receptor for the AMBRA1/CRL4 complex (Fig. 2A) we next examined whether AMBRA1/CRL4 can regulate Rab40c polyubiquitylation. To this end, we transfected 293T cells with FLAG-Rab40c, Myc-Ub, HA-AMBRA1, and HA-AMBRA1-DN (AMBRA mutant that does not bind Cullin4) individually or in various combinations (Fig. 3A). Lysates were then immunoprecipitated with anti-FLAG antibodies and blotted for Myc-Ub with anti-Myc antibodies. When Myc-Ub was co-transfected with FLAG-Rab40c in the presence of the proteasomal inhibitor MG132 high molecular weight species were detected, presuming polyubiquitinated Rab40c, which were significantly enhanced by co-transfecting HA-AMBRA1. Surprisingly, like wild type HA-AMBRA1, HA-AMBRA1-DN also stimulated Rab40c poly-ubiquitination (Fig. 3A&B), suggesting that AMBRA1/CRL4 ligase activity is not required for Rab40c ubiquitination. Although AMBRA1-DN no longer binds to Cullin4 it still binds to Rab40c (Fig. 1E), thus, we speculate that AMBRA1 binding to Rab40c promotes auto-ubiquitination of Rab40c by Rab40c/CRL5 complex. To test this hypothesis, we performed a similar ubiquitination assay as described above except that FLAG-Rab40c was replaced by FLAG-Rab40c-4A, a mutant that does not bind to Cullin5 and cannot be part of CRL5 complex [11]. As shown in the figure 3A&B, FLAG-Rab40c-4A ubiquitination was barely detected even with co-transfection with Myc-Ub, suggesting that Rab40c-4A lost self-catalyzed ubiquitination. Under these conditions, co-transfection with either AMBRA1 WT or AMBRA1-DN no longer had any effect on Rab40c-4A ubiquitylation (Fig. 3A&B). Taken together, these results demonstrate that AMBRA1 binding, but not AMBAR1/CRL4 ubiquitin ligase activity, is necessary for Rab40c ubiquitylation. Our data raises the possibility that AMBRA1 may enhance Rab40c self-ubiquitylation by regulating Rab40c interaction with Cullin5. To test this, we overexpressed FLAG-Rab40c individually or with HA-AMBRA1 or HA-AMBRA1-DN, followed by precipitation with anti-FLAG antibodies and immunoblotting for endogenous Cullin5. As shown in the figure 3C, FLAG-Rab40c can co-immunoprecipitate with endogenous Cullin5, as we reported previously [11]. Importantly, the amount of Cullin5 co-immunoprecipitatng with FLAG-Rab40c significantly increased when FLAG-Rab40c was co-transfected either with AMBRA1 or AMBRA1 DN (Fig. 3C). Therefore, we propose that AMBRA1 binding can increase Rab40c interaction with Cullin5, and results in Rab40c self-ubiquitination and activation independent of AMBRA1/CRL4 ubiquitin E3 ligase activity (Fig. 3D).

**Figure 3.**
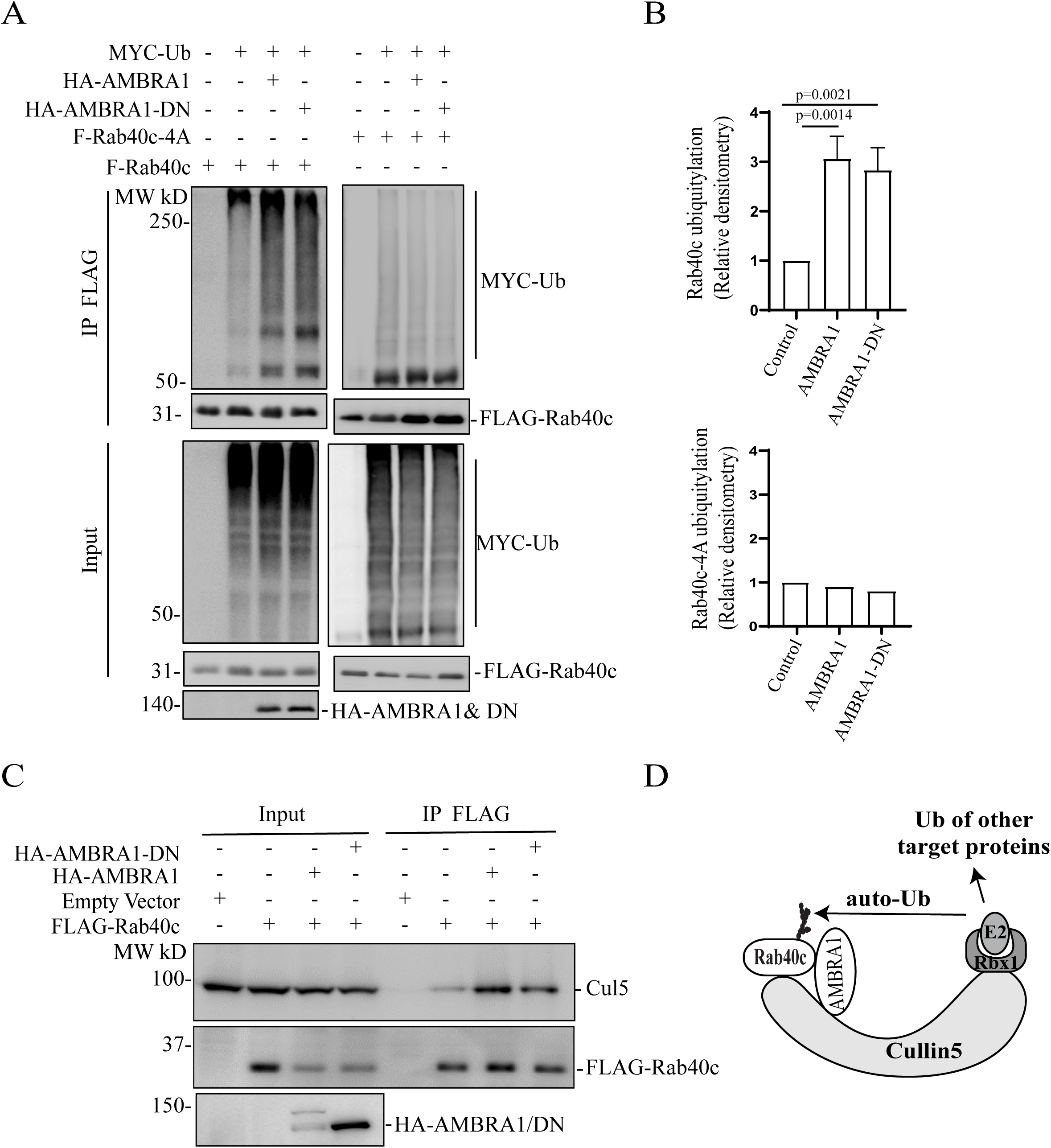
AMBRA1 regulates Rab40c binding to Cullin5 and auto-ubiquitylation. (A) *In vivo* Rab40c ubiquitylation assay. 293T cells were transfected with indicated plasmids and incubated for 48h. After being treated with 10μm MG132 for 6h, cells were harvested and immunoprecipitated with an anti-FLAG antibody, followed by Western blotting with anti-Myc, anti-FLAG, and anti-HA antibodies. Myc signal in anti-FLAG immunoprecipitate represents the extend of FLAG-Rab40c ubiquitylation. (B) Quantification of the levels of Rab40c polyubiquitylation shown in panel A. The data shown are the means and SEM derived from three different independent experiments. (C) 293T cells were co-transfected with indicated plasmids. FLAG-tagged Rab40c were then immunoprecipitated with anti-FLAG antibody and precipitates were immunoblotted with indicated antibodies. (D) A proposed model showing that AMBRA1 promotes Rab40c auto-ubiquitylation.

### AMBRA1 binding to Cullin4 is regulated by Rab40/CRL5-dependent ubiquitylation

Because the Rab40 subfamily of proteins are substrate receptors for CRL5 complex [5–7, 37], we next tested whether Rab40 can ubiquitylate and regulate AMBRA1. To this end, we transfected 293T cells with FLAG-AMBRA1, Myc-Ub, HA-Rab40c, or HA-Rab40c-4A (Rab40c mutant that does not bind Cullin5) individually or in various combinations (Fig. 4A). Lysates were then immunoprecipitated with anti-FLAG antibodies and blotted for Myc-Ub with anti-Myc antibodies. When Myc-Ub was co-transfected with FLAG-AMBRA1 in the presence of the proteasomal inhibitor MG132, high-molecular weight species were detected, presumably polyubiquitylated AMBRA1. AMBRA1 polyubiquitylation was enhanced by co-transfecting HA-Rab40c (Fig. 4A-B). Importantly, co-transfecting Cullin5-binding Rab40c mutant (HA-Rab40c-4A) decreased Rab40c-induced increase in AMBRA1 ubiquitylation. Note that HA-Rab40c-4A did not completely block AMBRA1 ubiquitylation (Fig. 4A-B), it suggested that AMBRA1 can also be ubiquitylated by other E3 ligases. Indeed, it was previously reported that AMBRA1 can self-ubiquitylate (as well as ubiquitylates other substrates) by forming CRL4 complex with Cullin4 and DDB1 (Fig. 4D) [22].

**Figure 4.**
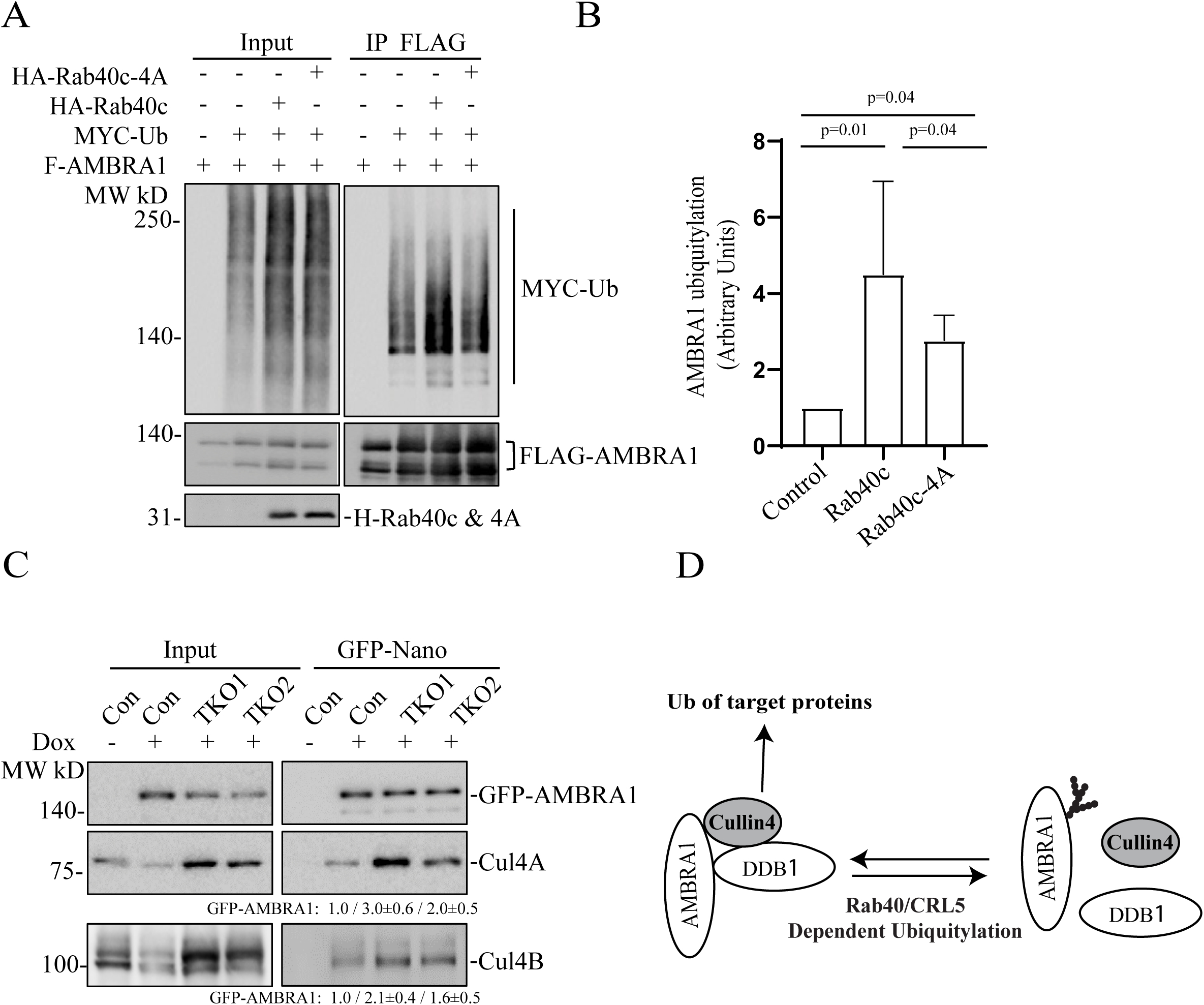
Rab40c/CRL5 ubiquitylates AMBRA1. (A) *In vivo* AMBRA1 ubiquitylation assay. 293T cells were transfected with indicated plasmids and incubated for 24h. After treatment with 100 nm Bafilomycin A1 overnight, cells were harvested and immunoprecipitated with an anti-FLAG antibody followed by Western blotting for either anti-Myc, anti-FLAG, or anti-HA. Myc signal in anti-FLAG immunoprecipitate represents the extent of FLAG-AMBRA1 ubiquitylation. (B) Quantification of FLAG-AMBRA1 ubiquitylation from panel A. The data shown represent the means and SEM derived from three independent experiments and are normalized against control. (C) MDA-MB-231 cells (control, TKO1 and TKO2) stably expressing dox-inducible GFP-AMBRA1 were incubated with 100 ng/ml doxycycline for 48h. GFP-AMBRA1 was then immunoprecipitated using anti-GFP-nanobody and immunoblotted with indicated antibodies. The quantification shown represents means and SEM derived from three independent experiments and normalized against control. (D) A proposed model showing that Rab40/CRL5 ubiquitylates AMBRA1 and inhibits AMBRA1/CRL4 complex formation.

Next, we set out to determine what are the functional consequences of AMBRA1 ubiquitylation by Rab40/CRL5. Given that AMBRA1 can interact with all members of Rab40 subfamily, we used Rab40a, Rab40b, and Rab40c triple-knockout MDA-MB-231 cells (TKO) [8] for the rest of the study. The best described AMBRA1 function is the formation of the AMBRA1/CRL4 complex that mediates polyubiquitylation and degradation of several proteins involved in cell proliferation and autophagy [23, 24, 27, 28, 30, 34, 38]. Thus, we next tested the effect of Rab40-TKO on the formation of the AMBRA1/CRL4 complex. To this end, we generated three different cell lines (control, TKO1, and TKO2) stably expressing dox-inducible GFP-AMBRA1. GFP-AMBRA1 was then immunoprecipitated using anti-GFP-nanobody and immunoblotted for the presence of Cullin4. As shown in the figure 4C, depletion of Rab40 increased Cullin4A and Cullin4B association with GFP-AMBRA1. Altogether, these results suggest that Rab40/CRL5-dependent ubiquitylation of AMBRA1 may lead to disassembly of the AMBRA1/CRL4 complex (Fig. 4D).

### AMBRA1 regulates transcription

Our data (see Figure 2) suggests that AMBRA1 affects Rab40c protein levels by regulating transcription of Rab40c mRNA. That raises an interesting possibility that AMBRA1 may have two distinct functions: to mediate protein ubiquitylation and proteasomal degradation as part of AMBRA1/CLR4 complex, and to regulate gene transcription. Indeed, a recent study reported that AMBRA1 is present in the nucleus where it appears to affect transcription [39]. What remains unclear is what genes are regulated by nuclear AMBRA1, and whether this regulation of transcription is dependent on AMBRA1/CRL4 ubiquitylation activity. Therefore, to identify the genes that are regulated by AMBRA1, we performed RNA sequencing (RNA-seq) analysis comparing control MDA-MB-231 cells with two different AMBRA1-KO MDA-MB-231 cell lines. Initial Principal Component analysis (PCA) demonstrated that AMBRA1-KO RNA samples were similar to each other, indicating the reproducible nature of their RNA content (Fig. 5A). Importantly, AMBRA1-KO RNA samples were well separated from control samples, suggesting that the AMBRA1-KO transcriptome is distinct from that of control cells (Fig. 5A). Compared with control cells, AMBRA1-KO led to downregulation of 194 genes and upregulation of 254 genes (KO/Control, Log2 > two-fold change, P < 0.05) (Fig. 5B). The gene ontology functional analysis revealed that many of these genes are involved in cell migration-related processes such as growth factor binding, extracellular matrix composition and organization, collagen binding, and cell-cell adhesion (Fig. 5C). Notably, we found that Rab40c mRNA is significantly increased in AMBRA1-KO cells (Fig. 5D), consistent with our previous observations (Fig. 2). To further confirm RNAseq results, we performed qPCR analysis on several selected mRNAs related to cell adhesion and migration, such as paxillin, MAP4K4, GRAMD1b, PXDN, and SNAI2 (Fig. 5D-E). Importantly, an increase in paxillin expression and a decrease in SNAI2 expression was also confirmed by Western blotting (Fig. 6A).

**Figure 5.**
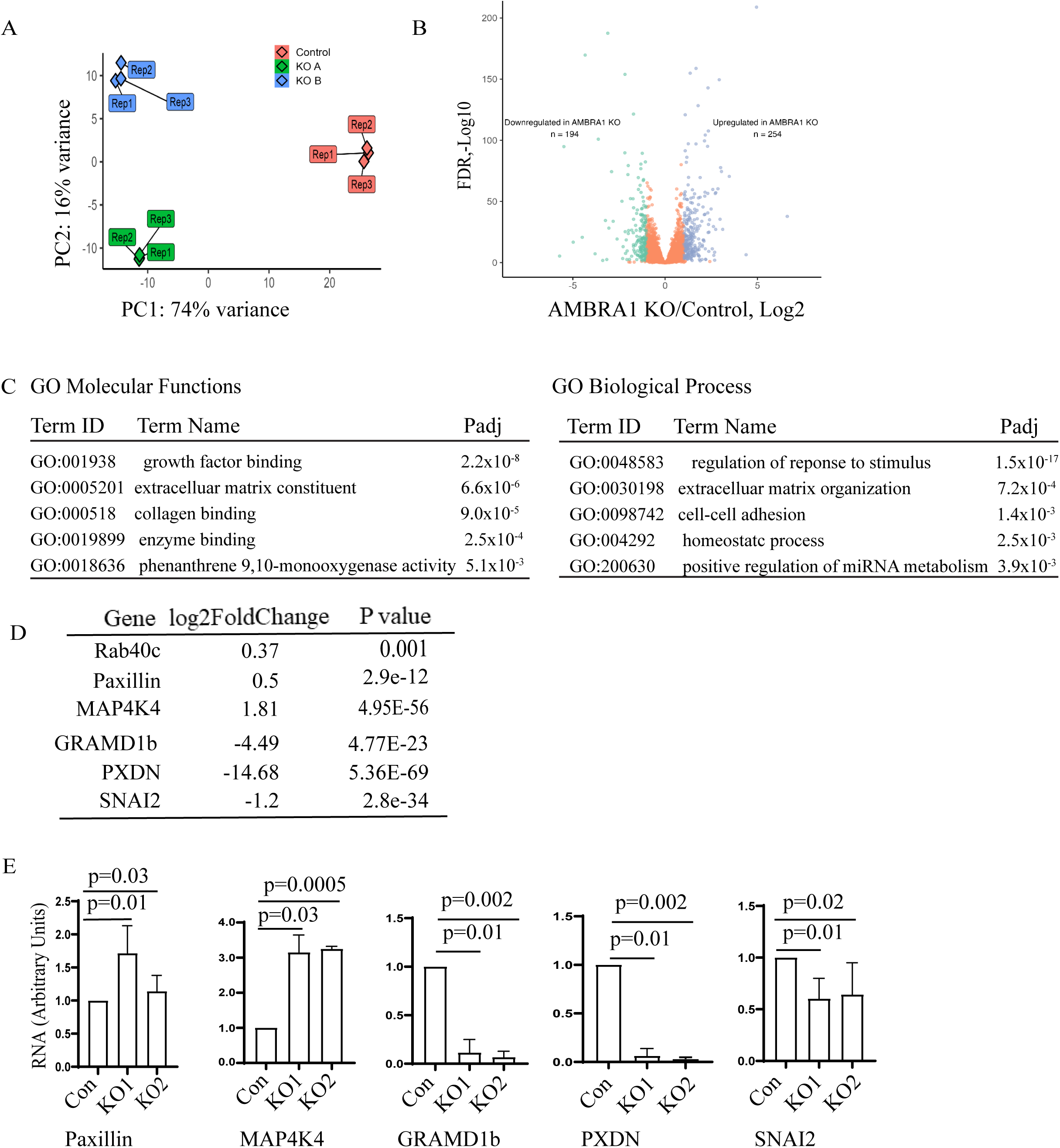
AMBRA1 regulates gene transcription. (A) Principal component analysis representing the differences between the control (3 reps) and the AMBRA1-KO (2×3 reps) RNAseq datasets. (B) Volcano plot showing the significantly downregulated (green) and significantly upregulated (blue) mRNAs identified in AMBRA1-KO cells. (C) GO molecular functions and biological process enrichment analysis of significantly changed mRNA (up- and down-regulated) identified by RNA-seq. (D-F) qPCR analysis of selected mRNAs that are either increased or decreased in AMBRA1-KO cells as indicted by RNA-seq. The means and SD were calculated from three independent experiments.

**Figure 6.**
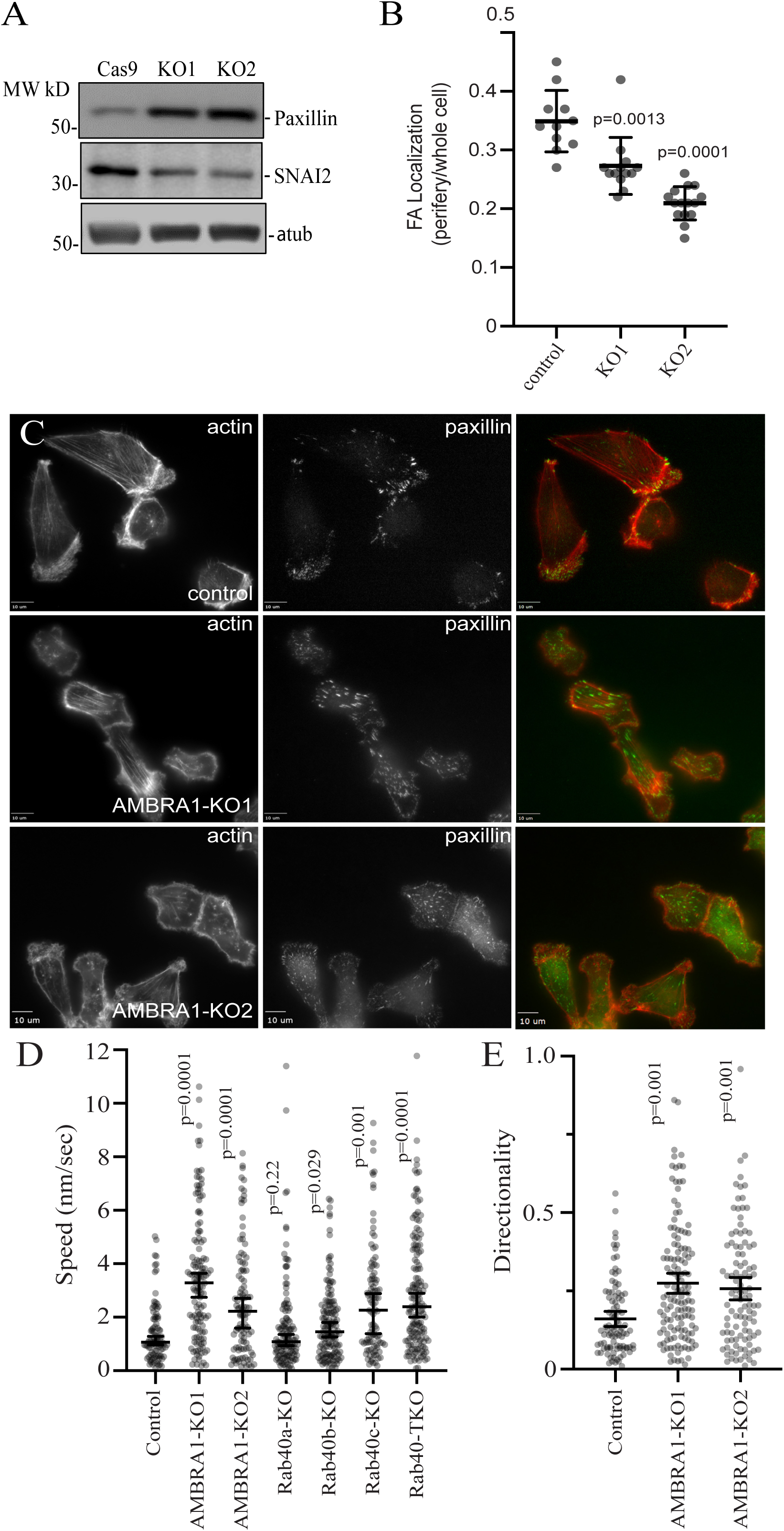
AMBRA1 regulates MDA-MB-231 cell migration. (A) To measure the levels of paxillin and SNAI2 cells lysates from control and AMBRA1-KO, cells were probed with anti-paxillin and anti-SNAI2 antibodies. Anti-α-tubulin antibodies were used as loading control. (B) Quantification of number of FAs at the cell periphery (within 4 μm from plasma membrane) for control and AMBRA1-KO cells. Data shown are means and SDs derived from three independent experiments. (C) Control, AMBRA1-KO1, and AMBRA1-KO2 MDA-MB-231 cells were plated on collagen-coated coverslips for 24 hours. Cells were then fixed and stained with phalloidin-Alexa Fluor 594 (red) and anti-paxillin antibodies (green). (D) Migration analysis of control, AMBRA1-KO, Rab40a-KO, Rab40b-KO, Rab40c-KO, or Rab40a/b/c TKO cells. For migration velocity quantification, 95-150 cells were randomly chosen for velocity tracking and velocity data (nm/sec) was extracted for each individual cell. The data shown are the means and SD derived from three biological replicates. Mann-Whittney’s test was performed to compare the velocities of cells from different cell lines. (E) Cell directionality was calculated as a ratio of the net displacement of a cell from its starting to final position compared with the total distance traveled from time-lapse analysis. Data shown represents the means and SEM derived from three different experiments. Mann-Whittney’s test was performed to compare the velocities of cells from different cell lines.

### AMBRA1 regulates cell migration

Our transcriptomic analysis has indicated that AMBRA1 may be involved in regulating cell migration. For example, RNA-seq, qPCR, and western blotting showed that AMBRA1 differently regulates paxillin and SNAI2 transcription and protein levels. Paxillin is a FA adapter which is involved in FA assembly/disassembly and signal transduction, thus, regulating cell adhesion and migration [40–43]. SNAI2 is a Snail family transcriptional repressor which is involved in regulating the epithelial-to-mesenchymal transition (EMT) and the migration of cancer cells [44–46]. We therefore investigated whether the cell motility and the structure of FAs changed in AMBRA1-KO cells. First, we assessed the number and distribution of FAs in control and AMBRA1-KO cells using an anti-paxillin antibody. As previously reported, in the control MDA-MB-231 cells paxillin-positive dot-like FAs were mostly present at the cell periphery, especially in leading-edge lamellipodia (Fig. 6B&C). In agreement with an increase of paxillin protein levels in AMBRA1-KO cells, an increase in the number and size of paxillin-positive FAs was observed in these cells (Fig. 6B&C). Strikingly, FAs did not accumulate at the periphery of the cell, but instead were scattered throughout the whole cells (Fig. 6B&C), suggesting AMBRA1 may regulate FA disassembly.

To directly test whether AMBRA1 regulates cell migration, we automatically tracked individual cell movements over time to quantify migration velocity. As shown in the figure 6D, compared with control cells, AMBRA1-KO cells exhibited increased individual cell migration and velocity. Importantly, MDA-MB-231 TKO cells (lacking Rab40a, Rab40b, and Rab40c) exhibited similar phenotypes, suggesting that Rab40 GTPases may regulate cell migration, in part, by binding and ubiquitylating AMBRA1. The change in FA size and distribution suggests that AMBRA1 may be involved in FA disassembly. Since stabilization of FAs is usually associated with an increase in cell adhesion and directionality, we next analyzed the directionality of control and AMBRA1-KO cells. As shown in Figure 6E, AMBRA1-KO cells did exhibit enhanced directionality during migration, supporting the hypothesis that Rab40-AMBRA1 pathway regulates cell migration by regulating FA dynamics.

### Transcriptional regulation by AMBRA1 does not require AMBRA1/CRL4-dependent ubiquitylation

Our data suggests that Rab40 GTPases bind and ubiquitylate AMBRA1. Furthermore, this Rab40-dependent ubiquitylation appears to regulate AMBRA1 function. Consequently, it would be expected that Rab40 knock-out and AMBRA1 knock-out would lead to similar changes in gene expression. To test this hypothesis, we performed qPCR analysis of selected transcripts that were affected in AMBRA1-KO cells (as determined by RNA-seq and qPCR). As shown in the figure 7A, knock-out of all three Rab40 isoforms (Rab40-TKO) led to a decrease in GRAMD1b, PXDN, and SNAI2 expression, as well as increase in MAP4K4 expression, the changes that were also observed in AMBRA1-KO cells (Fig. 5). Taken together, these data suggest that the Rab40-AMBRA1 pathway is involved in regulating gene transcription. SNAI2 is a key transcriptional factor for MDA-MB-231 migration and invasion, therefore, we selected it for further investigation.

**Figure 7.**
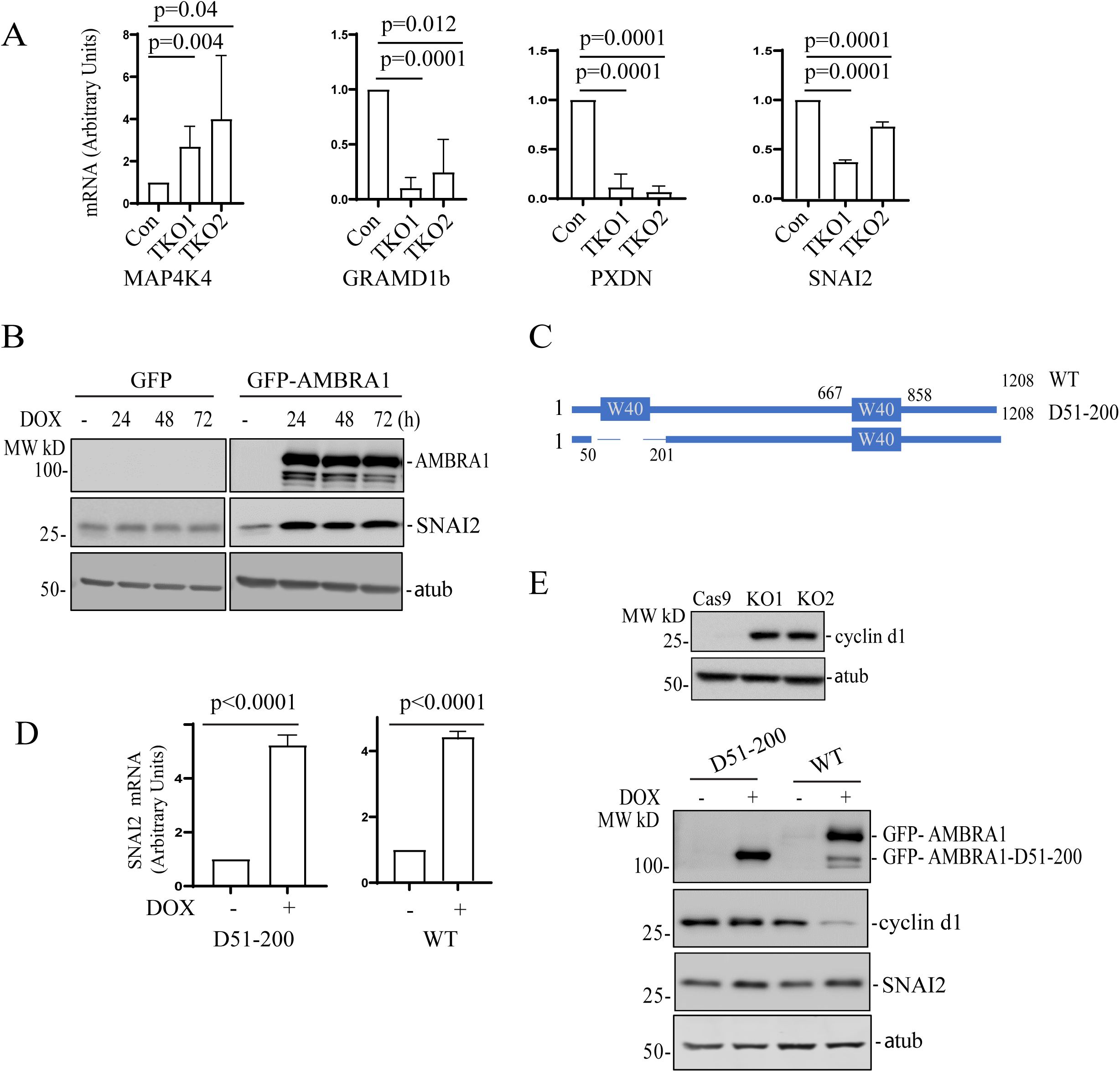
Transcriptional regulation by AMBAR1 is independent from AMBRA1/CR4 complex formation. (A) qRT-PCR analysis of selected mRNAs that were affected in AMBRA1-KO cells (see Figure 5E). RNAs were isolated from control and Rab40-TKO MDA-MB-231 cells. The data shown are the means and SD calculated from three independent experiments. (B) AMBRA1-KO MDA-MB-231 cells stably expressing Dox-inducible GFP or GFP-AMBRA1 were incubated with 100 ng/ml doxycycline for indicated time. Cell lysates were then immunoblotted with anti-SNAI2, anti-AMBRA1, and anti-α-tubulin (loading control) antibodies. (C) A schematic representation of AMBRA1 and AMBRA1 N-WD40 deletion mutant. (D) qRT-PCR analysis of the levels of SNAI2 mRNA in AMBRA1-KO MDA-MB-231 cells stably expressing either dox-inducible GFP-AMBRA1-Δ51−200 or dox-inducible GFP-AMBRA1. The means and SD were calculated from three independent experimentals. (E) Top panels: Immunoblotting of cell lysates from control and AMBRA1-KO cells with anti-cyclin d1 and anti-α-tubulin (loading control) antibodies. Lower panels: AMBRA1-KO MDA-MB-231 cells stably expressing dox-inducible GFP-AMBRA1-Δ51−200 or GFP-AMBRA1 were incubated with 100 ng/ml doxycycline for 48h. Cell lysates were immunoblotted with anti-AMBRA1, anti-SNAI2, anti-cyclin d1, and anti-α-tubulin antibodies.

While our data suggests that AMBRA1 regulates gene transcription, the underlying mechanism governing this AMBRA1 function remains unclear. One possibility is that this function is dependent on Cullin4A/B, since Cullin4A/B were both reported to localize to the nucleus, where they appear to mediate multiple nucleus-related functions, including DNA replication/repair and chromatin remodeling [47–50]. Another possibility is that the cytosolic AMBRA1/CRL4 complex ubiquitylates selected transcription regulators, thus, affecting their protein levels and/or their translocation to nucleus. AMBRA1 contains a N-terminal WD40 domain which is responsible for binding Cullin4A/B and DDB1 E3 ligase to form a AMBRA1/CRL4 complex. To test whether AMBRA1 transcription regulatory function is dependent on CRL4 binding, we generated two AMBRA1-KO cell lines stably expressing a dox-inducible GFP-tagged wild-type AMBRA1 (WT) or N-terminal WD40 deletion mutant (D51-200) that cannot bind to Cullin4 (Fig. 7B-C) [14, 33]. To confirm that the AMBRA1(D51-200) mutant cannot mediate proteasomal degradation, we immunoblotted cell lysates using anti-cyclin D1 antibody. Cyclin D1 is a known target of AMBRA1/CRL4-dependent ubiquitylation and degradation [15, 27, 29, 30, 51]. Consistent with previous reports, AMBRA1 knock out leads to an increase in cyclin D1 protein levels and results in a delay in entering the G2/M phase (Fig. S2). This AMBRA1-KO induced increase in cyclin D1 proteins levels can be eliminated by over expressing WT AMBRA1 but not the AMBRA1(D51-200) mutant (Fig. 7C), demonstrating that AMBRA1/CRL4 ubiquitylation activity is needed to regulate cyclin D1 levels. Next, we tested whether AMBRA1/CRL4 ubiquitylation activity is required for an increase in SNAI2 mRNA. To this end, we measured SNAI2 mRNA levels in MDA-MB-231 cells by qPCR. Induction of GFP-AMBRA1 expression in AMBRA1-KO cells increased SNAI2 mRNA levels which is consistent with our RNA-seq result, suggesting AMBRA1 is a positive regulator for SNAI2 expression (Fig. 7D). Intriguingly, SNAI2 mRNA also significantly increased when AMBRA1(D51-200) expression was induced, suggesting that AMBRA1 activates SNAI2 transcription independent of AMBRA1/CRL4 complex formation (Fig. 7C). Lastly, SNAI2 expression stimulated by AMBRA1 and AMBRA1(D51-200) was also confirmed by western blotting (Fig. 7E). All these data strongly suggest that in addition to its function as a component of the E3 ligase complex, AMBRA1 may also function as a transcriptional regulator, which is dependent on its interaction with Rab40 GTPases.

### The roles of different AMBRA1 splice-isoforms in regulation of transcription

Our data so far identified a novel function for AMBRA1 as a transcriptional regulator that is not dependent on its ability to bind Cullin4 and CRL4 mediated protein ubiquitylation. What remains unclear is whether AMBRA1 directly regulates transcription by translocating to the nucleus, or whether it acts as a cytosolic scaffolding protein that indirectly affects transcription by sequestering other transcriptional regulators in cytosol. Importantly, AMBRA1 exists as several different splice isoforms (Fig. S3A). The main difference between these isoforms is the presence or absence of two insertions, one at aa601 (isoform 1, ISO1) and a second at aa255 (isoform 5, ISO5). We used isoform 2 for all overexpression analyses shown in this study. Thus, we next asked whether these different isoforms have a differential function in regulating protein ubiquitylation/degradation and transcription. To this end, we generated two additional MDA-MB-231 AMBRA1-KO cell lines stably expressing dox-inducible isoforms 1 and 5 of AMBRA1 (Fig. S3B). These cell lines were then tested for their ability to target cyclin D1 for proteasomal degradation. As shown in the figure S3B, isoform 1 and isoform 5 could both induce cyclin D1 degradation in a manner like isoform 2 (Fig. 7C). Next, we used qPCR to test whether isoforms 1 and/or 5 can increase SNAI2 expression. As shown in the Figure S3C, all tested AMBRA1 isoforms can stimulate SNAI2 transcription.

While our data demonstrates that, when overexpressed, all AMBRA1 splice isoforms can induce SNAI2 expression and cyclin D1 proteasomal degradation, we wondered whether there are differences in nuclear import between these AMBRA1 isoforms. To this end we fractionated isoform 1, isoform 2, and isoform 5 expressing cells into nuclear (N) and cytosolic (C) fractions and compared the distribution of different AMBRA1 isoforms between cytosol and nucleus. Interestingly, while all isoforms could be detected in the nucleus, isoform 5 appears to be targeted to nucleus more efficiently than isoforms 1 and 2 (Fig. S4A). To further confirm AMBRA1 nuclear localization, we next used immunofluorescence microcopy to analyze subcellular distribution of all three AMBRA1 isoforms. As shown in the figure S4B, AMBRA1 isoforms 1 and 2 were predominately present in cytosol. In contrast, AMBRA1 isoform 5 could clearly be observed in the nucleus and perinuclear organelles. Thus, while further research is needed to better define the roles of different AMBRA1 splice isoforms, our data suggests that isoform 5 may have distinct subcellular localization and function as compared to other AMBRA1 isoforms.

## DISCUSSION

The Rab40/CRL5 protein complex has recently emerged as a unique Rab40-dependent complex that plays an important role in regulating spatiotemporal dynamics of protein ubiquitylation during cell migration. Intriguingly, in many cases this ubiquitylation does not induce target protein degradation, but rather regulates their localization and activity [8, 11, 12]. In this study, we identified AMBRA1 as a novel Rab40/CRL5 substrate protein and have shown that Rab40/CRL5 E3 ligase mediates non-proteolytic polyubiquitylation of AMBRA1. Surprisingly, we found that AMBRA1 ubiquitylated by Rab40/CRL5 appears to function as a transcriptional regulator and that AMBRA1 affects transcription independent of its ability to bind Cullin4 and mediate protein ubiquitylation. Thus, our findings expand the current understanding of molecular functions of AMBRA1 and suggest that AMBRA1 has at least two distinct functions. The first one is the canonical function of regulating protein polyubiquitylation and degradation during activation of autophagy. The second one is transcriptional regulation of selected subsets of mRNAs, the process that is independent from AMBRA1/CRL4 enzymatic activity.

AMBRA1 is an intrinsically disordered protein which has been shown to bind numerous other proteins and perform many functions. Although we used multiple approaches to establish that Rab40 subfamily GTPases as novel binding partners of AMBRA1, it remains unclear if their interaction is direct or mediated by other proteins. We did show that two distinct motifs, one at the C-terminus and one at the N-terminus, are required for AMBRA1 interaction with Rab40. Both of these Rab40-interacting motifs are located in close proximity to split-WD40 domains. Since it was shown that these split-WD40 domains in AMBRA1 form a fully functional WD40 domain, it is likely that both Rab40-interacting motifs are brought in close proximity to form a Rab40-binding interface, although further experiments will be needed to demonstrate that. While Rab40 binds both AMBRA1 and CRL5, we could not detect AMBRA1 binding (via Rab40) to CRL5. One possibility is that AMBRA1 is a Rab40/CRL5 substrate and AMBRA1 ubiquitylation would likely cause its rapid dissociation from the Rab40C/Cul5 complex. Consistent with the idea, we found that the Rab40c-4A mutant (does not bind Cullin5, thus enzymatically inactive) binds to AMBRA1 more strongly than wild-type Rab40c.

AMBRA1 is a substrate receptor for CRL4 while Rab40s and acts as the adaptor proteins for CRL5, therefore, their interaction may result in ubiquitylation of each other. In fact, we observed that overexpression of AMBRA1 can stimulate Rab40c ubiquitylation. However, AMBRA1-ΔWD40, a mutant unable to bind DDB1, can also stimulate Rab40c ubiquitylation, suggesting Rab40c is not a substrate of CRL4 but rather that their interaction stimulates Rab40c self-ubiquitylation. In support of this, we found that AMBRA1 lost its ability to stimulate ubiquitylation of Rab40c-4A, which cannot bind to Cullin5 and form the Rab40/CRL5 complex [11]. Since AMBRA1 appears to increase Rab40c binding to Cullin5, AMBRA1 may function as a positive regulator of the Rab40/CRL5 complex assembly, differing from two recent reports showing that AMBRA1 represses CRL5 activity by binding Elongin B or by targeting Elongin C for degradation [22, 23]. The reason for these different effects of AMBRA1 on CRL5 activity is unclear, but we speculate that AMBRA1 may differentially modulate CRL5 signaling during different cellular processes such as autophagy (binding Elongin B), inflammation (degrading Elongin C), or cell migration (promoting Rab40/CRL5 assembly). Interestingly, we found that AMBRA1 is a substrate of Rab40c/CRL5-dependent ubiquitylation. Previous studies have shown that AMBRA1 undergoes post-translational modifications, including Cullin4-dependent self-ubiquitylation of AMBRA1, or RNF2 mediated AMBRA1 polyubiquitination, leading to proteasomal degradation [22, 52, 53]. In this study we suggest that Rab40/CRL5-mediated AMBRA1 ubiquitylation does not lead to its degradation, but instead appears to mediate the disassembly of the AMBRA1/CRL4 complex.

Although Rab40c is not a substrate of AMBRA1/CRL4-dependent ubiquitylation and proteasomal degradation, Rab40c protein levels were significantly increased in AMBRA1 KO cells. Interestingly, we demonstrate that Rab40c mRNA levels increase in AMBRA1 KO cells, suggesting that AMBRA1 may regulate (directly or indirectly) Rab40c transcription. In reminiscence of a recent study in which AMBRA1 was found to bind with multiple classes of proteins in the nucleus to regulate gene transcription, we set to further dissect the role of AMBRA1 in regulating transcription and to identify the global gene profiles affected by AMBRA1 depletion. The RNA-seq results confirmed that Rab40c mRNA did indeed increase in AMBRA1 KO cells. Importantly, AMBRA1 depletion led to changes in transcription of multiple genes, suggesting that AMBRA1 may be a transcriptional regulator of these subsets of genes. How AMBRA1 regulates gene transcription remains unclear and will be a subject of further studies. Since AMBRA1 does not have clearly identifiable DNA-binding domain, it is unlikely that AMBRA1 directly binds to the promotor or enhancer regions. One possibility is that AMBRA1 is recruited to specific promoters by binding chromatin-modifying proteins. Indeed, it was previously suggested that AMBRA1 can regulate histone methylation [39]. Another possibility is that AMBRA1 regulates the activity of some transcriptional activators or repressors by either directly binding to them (and acting as scaffolding factor) or mediating their ubiquitylation. For example, a recent study showed that AMBRA1/CRL4 mediates non-proteolytic polyubiquitylation of Smad4 to enhance its transcriptional functions [54]. Surprisingly, we have shown that AMBRA1-dependent transcriptional regulation does not require AMBRA1/CRL4 E3 ligase complex. That raises an intriguing possibility that AMBRA1 may have a non-canonical function in regulating gene transcription independent of CRL4 mediated ubiquitylation. However, further studies will be needed to confirm that and to dissect the molecular machinery governing AMBRA1-dependent regulation of transcription.

It is well-established that protein ubiquitylation plays a crucial role in regulating gene transcription [55–57]. Non-proteolytic polyubiquitylation of AMBRA1 by Rab40/CRL5 may affect AMBRA1-mediated transcription. Consistent with this idea, we found that transcription of some selected genes is affected in AMBRA1 KO and Rab40 TKO cells, suggesting that Rab40 and AMBRA1 co-regulate the same subset of genes. In this regard, Rab40/CRL5 establishes a negative feedback mechanism to regulate itself by ubiquitylating AMBRA1, which then represses (directly or indirectly) Rab40 transcription. Intriguingly, while we show that AMBRA1 regulates transcription, it does not have a canonical nuclear localization signal (NLS). Previous studies suggested that CRL4s are localized in the nucleus, especially for CRL4B, which contains a NLS in its N-terminus [47]. However, we found that AMBRA1-dependent transcriptional regulation does not require binding to CRL4s. Thus, AMBRA1 targeting into the nucleus is likely dependent on other factors. It is important to note that we cannot rule out the possibility that AMBRA1 may indirectly regulate transcription by binding and sequestering transcriptional regulators in the cytosol.

What are the genes regulated by AMBRA1? Gene ontology enrichment analysis of differentially expressed genes revealed that many affected genes are involved in extracellular matrix composition and remodeling, as well as cell-substrate adhesion. Indeed, previous evidence indicates that AMBRA1 can be recruited to FAs where it controls the levels of active Src and FAK [32]. We found that paxillin, a FA adapter protein, mRNA, and protein levels were significantly increased in AMBRA1 KO cells. In contrast, the mRNA of SNAI2, a zinc-finger transcription factor which is involved in epithelial-mesenchymal transitions, was significantly decreased. All these findings suggest that regulation of gene transcription by AMBRA1may affect cell migration. Consistent with this hypothesis, we observed that AMBRA1 depletion affected the number and position of FAs, as well as the speed and directionality of cell migration. Importantly, we observed MDA-MB-231 TKO cells (lacking Rab40a, Rab40b, and Rab40c) exhibited similar effects on cell migration, further supporting the hypothesis that Rab40-dependent ubiquitylation enhances AMBRA1 transcriptional effects.

AMBRA1 is a multifunctional protein that is involved in many cellular processes, including autophagy, proliferation, apoptosis, transcription, and cancer drug resistance [17, 34, 58–62]. Here we examined the potential roles of AMBRA1 in regulating transcription and cell migration. Furthermore, we show that Rab40/CRL5-dependent ubiquitylation of AMBRA1 appears to be required for its transcriptional function. However, many questions remain. We do not know what AMBRA1 amino acid residues are ubiquitylated by Rab40/CRL5 and what type of ubiquitylation (Lys63 or Lys48) is mediated by Rab40/CRL5. It is also unclear whether Rab40/CRL5-dependent ubiquitylation also affects canonical AMBRA1 function, such as regulating autophagy. Finally, we are only beginning to understand how AMBRA1 regulates transcription, and what the functions are of different AMBRA1 splice-isoforms. Further studies will be needed to address all these questions.

## MATERIALS AND METHODS

### Cell Culture and Transfection

All cell lines were cultured as described previously [11, 12, 63]. Briefly, human embryonic kidney (HEK) 293T cells were grown in complete Dulbecco’s modified Eagle medium (DMEM supplemented with 10% fetal bovine serum and 100 μg/ml of penicillin and streptomycin) at 37 °C in a 5% CO2 atmosphere. MDA-MB-231cells were grown in complete DMEM supplemented with 1ug/ml human recombinant insulin, 1% non-essential amino acids, and 1% sodium pyruvate. Cell lines were routinely tested for mycoplasma. All cell lines used in this study were authenticated and are in accordance with American Type Culture Collection standards. 293T cells were grown to 60–70% confluence and transfected using the standard calcium phosphate precipitation method [63, 64]. Typically, 10 μg of plasmid was used for a single-gene transfection of a 100-mm dish of cells, with up to 30 μg of plasmids used for co-transfection of three plasmids. MDA-MB-231 cells were grown to 80-90% confluence and transfected using JetPRIME (polyplus). Lipofectamine RNAiMAX (Invitrogen) was used for transfection of siRNAs both in 293T and MDA-MB-231 cells.

### Antibodies and reagents

The following antibodies were used in this study: anti-FLAG (clone M2, WB 1:1,000, Sigma), paxillin (IF 1:500 Transduction labs). Anti-GAPDH (WB 1:5,000) and anti-GFP (GF28R, WB 1:2,000) were purchased from UBPBio. Rabbit anti-AMBRA1(WB 1:1000), Rabbit anti-Cul4A (WB 1:2,000), and Rabbit anti-Cul4B (WB 1:2,000) were purchased from Proteintech. Mouse anti-c-MYC (9E10, WB 1:1,000), mouse anti-HA (WB 1:500, SC F-7), mouse anti–α-tubulin (WB 1:3,000), anti-Rab40c (WB 1:500, H-8), cul-5 (WB 1:500, H-300), and mouse anti-AMBAR1(G-6, 1:500) were purchased from Santa Cruz Biotechnology. Rabbit anti-cyclin d1(WB 1:1,000), Rabbit anti-SNAI2 (C19G7, WB 1:1,000), Rabbit anti-HP1 (WB, 1:1,000), and Rabbit anti-H2AK119U (WB 1:2,000) were purchased from cell signaling technology. TRIzol, Puromycin, and Doxycycline were purchased from Thermo Fisher Scientific. MG132 and Bafilomycin A1 were purchased from Selleckchem. Complete protease inhibitor cocktail and phosphatase inhibitor cocktails were purchased from Roche.

### Mammalian Expression Constructs

Human Rab4A, AL, B, C, FLAG-Rab40c-4A mutant, and Myc-Ub plasmids were described previously [11]. pcDNA4-AMBRA1-3xFLAG (174157), pcDNA4-deltaH-AMBRA1-3xFLAG (#174158), and pCW-Cas9 (#50661) were obtained from Addgene. AMBRA1 isoform1 was purchased from GeneCopoeia. AMBRA1 isoform 5 was cloned by PCR. GFP-AMBRA1 and GFP-AMBRA1 □51-200, FLAG-AMBRA1 deletion mutants were constructed by PCR, followed by subcloning into the pRK7 or pCW vector containing an N-terminal GFP or FLAG tag. All plasmids were validated by DNA sequencing.

### Immunoprecipitation and Western blot analysis

For non-denaturing immunoprecipitation, cells in a 100-mm dish were harvested and lysed on ice in buffer containing 20 mM Tris-HCl, pH 7.4, 150 mM NaCl, 2 mM EDTA, 1% Triton X-100, 10% glycerol with protease inhibitor, and phosphatase inhibitor cocktails. After clearing lysates by centrifugation, supernatants were incubated with 2μg of an appropriate antibody or control IgG for 4 h at 4°C, then supplemented with 50 μl protein G beads. After overnight incubation, the protein G beads were pelleted by centrifugation and washed three times with 1 ml of lysis buffer plus 0.5 M NaCl. Bound proteins were eluted in 50 μl 1× SDS sample buffer. For denaturing immunoprecipitation, cells in a 100-mm dish were lysed in 1 ml cell lysis buffer plus 1% SDS. Cell lysates were collected and then heated at 95°C for 10 min. After centrifugation, supernatants were diluted with the cell lysis buffer to reduce SDS concentration to 0.1%. The immunoprecipitation assay was performed as described above, except that 5 μg anti-FLAG M2 antibody was used in each reaction. Eluates were then resolved in SDS-PAGE and transferred to nitrocellulose membranes for immunoblotting assays. Immunoblotting images were captured using a ChemiDoc MP Imaging system (Bio-Rad).

### MDA-MB-231 CRISPR/Cas9 KO cell lines and genotyping

MDA-MB-231 cells stably expressing Tet-inducible Cas9 (Horizon Discovery Edit-R lentiviral Cas9) were grown in a 12-well plate to ∼75% confluency and then treated with 1 µg/ml doxycycline (dox) for 24 h to induce Cas9 expression. After 24 h, cells were cotransfected with crRNA:tracrRNA mix using DharmaFECT Duo transfection reagent as described by the Horizon Discovery DharmaFECT Duo protocol. crRNAs for AMBRA1 targeting are 5’-TACCATTACTGATTTCAGGG-3’ (exon 12, isoform 2) and 5’-ACTGACATGTCTCCGCTGGT-3’ (exon16 isoform 2). Cells were split 24 h after transfection and seeded for single colonies and then were screened by WB followed by PCR cloning and genotyping. Rab40a, b, or triple knockout lines have been described previously [8]. For each knock-out line, two different clonal lines were used for experiments.

### Quantitative PCR (qPCR)

Total RNA was extracted using TRIzol (Invitrogen) according to the manufacturer’s protocol. Reverse transcription to cDNA was performed with SuperScript IV (Invitrogen) using oligo(dT) primers. qPCR was performed using iTaq SYBR Green qPCR Master Mix on Applied Biosystems ViiA7 Real Time PCR System. The qPCR amplification conditions were 50°C (2 min), 95°C (10 min), 40 cycles at 95°C (15 s), and 60°C (1 min). Targets were normalized to GAPDH. The following primers used for qPCR were from PrimerBank (https://pga.mgh.harvard.edu/primerbank/): RAB40C forward 5′-GGCCCAACCGAGTGTTCAG-3′ and reverse 5′-GGACTTGGACCTCTTGAGGC-3’; AMBRA1 forward 5′-CTCTTCCTCAGACAACCAGGGT-3′ and reverse 5′-TCCAAGCGAAGGTGCAGACATC-3’; paxillin forward 5′-ACAGTCGCCAAAGGAGTCTG-3′ and reverse 5′-GGGGCCGTTGCAGTAGTAG-3’; MAP4K4 forward 5′-GGAACACACTCAAAGAAGACTGG-3′ and reverse 5′-GTGCCTATGAACGTATTTCTCCG-3′; GRAMD1b forward 5′-GCTATGGGAACGAATTGGGC-3′ and reverse 5′-CTGCTCTTGGATGAGCTGTCA-3′; SNAIL2 forward 5′-TGTGACAAGGAATATGTGAGCC-3′ and reverse 5′-TGAGCCCTCAGATTTGACCTG-3′; PXDN forward 5′-AATCAGAGAGATCCAACCTGGG-3′ and reverse 5′-AATGCTCCACTAGGTATCCTCTT-3′; GAPDH forward 5′-CTGGGCTACACTGAGCACC-3′ and reverse 5′-AAGTGGTCGTTGAGGGCAATG-3′.

### Cellular fractionation

Cytoplasmic and nuclear/cytoskeletal fractions were isolated using the Cell Fractionation Kit from Cell Signaling Technologies (#9038) according to the manufacturer’s instructions. Briefly, a 10cm dish of MDA-MB-231 cells at 100% confluency were trypsinized and washed in PBS. The cell pellets were resuspended in 500µl CIB, vortexed for 5sec, incubated on ice for 5min, and centrifuged for 5 min at 500 x g. The supernatant was saved as the cytoplasmic fraction. The pellet was resuspended in 500µl MIB, incubated on ice for 5min, and centrifuged for 5 min at 8,000 x g to remove the supernatant (the membrane and organelle fraction). The pellet was resuspended directly in 50µl 1X SDS as the cytoskeletal and nuclear fraction.

### *In vivo* ubiquitination Assay

*In vivo* ubiquitination assay was performed as described previously[8, 12, 63]. Briefly, 293T cells (∼70% confluency) were transfected with various combinations of plasmids including MYC-Ub. After 24 hrs, cells were treated with 10 μM MG132 for 6 h or 100nM Bafilomycin-A1 (Selleckchem S1413) overnight. Then, cells were lysed in 1% SDS for denaturing immunoprecipitation as described above. Bound proteins were eluted in 50µl 1X SDS sample buffer. Eluates (20 µl) were resolved via SDS-PAGE and transferred to nitrocellulose membranes for immunoblotting.

### siRNA knockdown

AMBRA1 siRNAs (SASI_Hs01_00116731 and SASI_Hs01_00116732) and mission siRNA universal negative control (SIC001; Sigma) were purchased from Sigma. siRNAs were transfected using Lipofectamine RNAiMAX (Invitrogen) according to manufacturer protocol.

### RNA-seq

RNA sequencing was performed by National Jewish Health Genomics Facility. Briefly, Total RNA was isolated using TRIzol (Invitrogen) according to the manufacturer’s protocol. RNA sequencing libraries were prepared according to the KAPA mRNA HyperPrep Library Build user guide. mRNA from 50 ng of total RNA was isolated using polyA, oligo-dT magnetic beads. The isolated mRNA was then subject to enzymatic fragmentation, resulting in 200-300 bp fragments. The resulting RNA fragments underwent first and second strand cDNA synthesis. Unique KAPA Dual-Indexes were then ligated to the cDNA. The ligated product was then PCR amplified for 13 cycles. The resulting libraries were quantified using the Qubit HSDNA assay and the TapeStation HSDNA 1000 assay. Equal molar concentrations were pooled, diluted, and sequenced using a NovaSeq 6000.

### Flow cytometry

Flow cytometry was used for cell cycle analysis which was conducted by CU Cancer Center Flow Cytometry Shared Resource. MDA-MD-231 cas9 control and two AMBRA1 KO cell lines were stained by propidium iodide (PI) as described previously [65]. Cells are analyzed using a Beckman Coulter Gallios flow cytometer. Doublets are excluded from the analysis using the peak vs. integral gating method. ModFit LT software (Verity Software House, Topsham, ME) is used for cell cycle analysis.

### Cell migration assays

For single cell migration assay, time-lapse imaging was performed using OLYMPUS IX83 inverted confocal microscope, with a brightfield 4X magnification objective equipped with a humidified chamber and temperature-controlled stage top. 35mm glass bottom dishes were coated with Fibronectin (catalogue number: F4759-1mg from Sigma Aldrich) and allowed to set for 1 hour under UV at room temperature. Once dried the cells were plated and allowed to attach for 24 hours. All time-lapses were taken at 20 minute intervals each, 36 frames were taken, resulting in a total time-lapse of ∼12 h. For cell migration analysis, cells were manually tracked using the Manual Tracking software Excellence pro. Generated data was acquired from this software, such as speed, which was used to calculate velocity. Careful effort was made to select the geometric center of the cell to focus on the nucleus when manually tracking. Three independent biological replicates were performed for each cell line. Statistical analysis (non-parametric student t-test) and Mann-Whittney’s test was performed on the averages of the three biological replicates and processed using Graph Prism.

### Immunofluorescent microscopy and Image analysis

MDA-MB-231 cells were seeded onto collagen-coated glass coverslips and grown in full growth media unless otherwise noted for at least 24 h. Cells were washed with PBS and fixed in 4% paraformaldehyde for 15 min. Samples were then washed three times in PBS then incubated in blocking serum (1× PBS, 5% normal donkey serum, and 0.3% Triton X-100) for 1 h at room temperature. Primary antibodies were then diluted at 1:100 in dilution buffer (1× PBS, 1% BSA, and 0.3% Triton X-100) and incubated for 1 hour at room temperature. Samples were then washed three times with PBS and incubated with fluorophore-conjugated secondary antibodies (1:100 in dilution buffer) for 30 min at room temperature. Cells were then washed three times in PBS and mounted onto glass slides. Cells were then imaged on either an inverted Zeiss Axiovert 200M deconvolution microscope with a 63× oil-immersion lens and Sensicam QE charge-coupled device camera, or a Nikon A1R. Z-stack images were taken at a step size of 100–500 nm.

Analysis of FA number and size FA quantification performed in ImageJ and was adapted from Horzum et al. (2014). For each experimental replicate, cells were analyzed from at least five randomly chosen fields. In total, 15-20 cells were analyzed for each experimental replicate. Only cells that did not make any contact with surrounding cells were analyzed. The same exposure was used for all images for each experimental replicate. Maximum intensity projections for relevant z-planes were created, and images were loaded into ImageJ. Background was minimized using the Subtract Background and EXP tools, images were filtered using the Log3D/ Mexican Hat plugin, and thresholds were then applied manually (method = default). Individual cells were defined by hand, and FAs were determined with the “Analyze Particle” command. Resulting particle outlines were then compared with the original image to ensure fidelity of the analysis. To analyze the number of FAs in cell periphery, the 4 μm wide area-of-interest around the cell was selected at the plasma membrane. The FAs in that area were then counted and the ratio of FAs in periphery/whole cell was calculated for each individual cell.

### Statistical analysis

Statistical analysis for all experiments was determined using GraphPad Prism Software (GraphPad). A two-tailed Student’s t test was used to determine statistical significance unless otherwise noted. Data were collected from at least three independent experiments unless otherwise noted. In all cases, P ≤ 0.05 was regarded as significant. Error bars represent standard errors unless otherwise noted. For all IF experiments, at least five randomly chosen image fields per condition were used for each experimental replicate. In total, 15-20 cells were analyzed for each experimental replicate, and each individual cell was treated as technical replicate. Statistical analysis was performed on means calculated from individual cells for each experimental replicate.

## ACKOWLEDGEMENTS

We would like to thank the University of Colorado Anschutz Medical Campus Genomics Facility and the RNA Biosciences Initiative (RBI) for analyzing the RNAseq data. We are grateful to Migle Prekeryte for critical reading and editing of the manuscript. This work was funded by the National Institutes of Health grant R01 GM122768 to R.P, grant S-MIP-22-60 from Research Council of Lithuania (to R.P.), and Diversity Supplement GM143774-02S1 to KV. The authors declare no competing financial interests.

## SUPPLEMENTAL FIGURE LEGENDS

**Figure S1.**
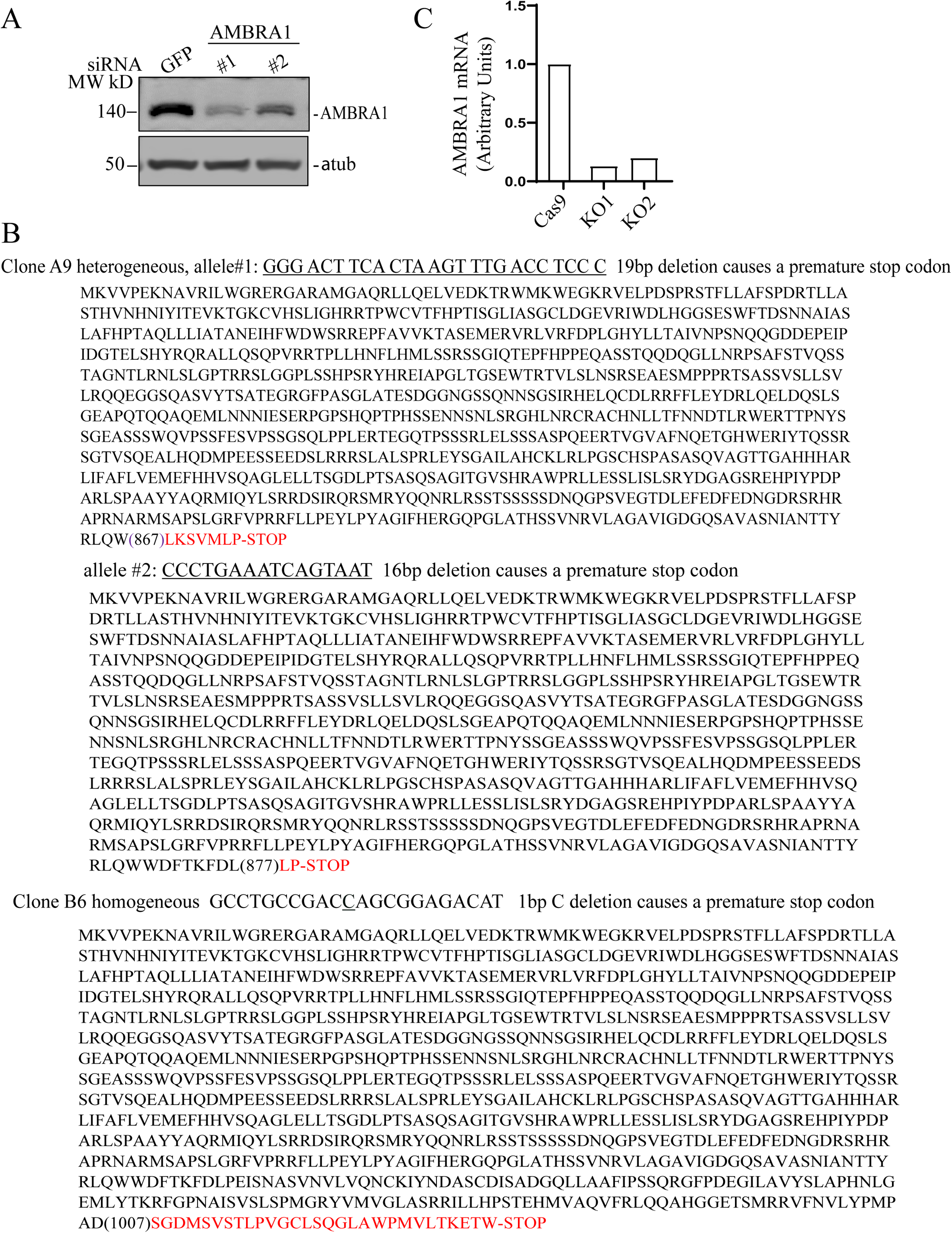
(A) To test the specificity of the anti-AMBRA1 antibody, MDA-MB-231 cells were transfected with non-targeting control siRNA or siRNA targeting AMBRA1. Cell lysates were then blotted with anti-AMBRA1 or anti-α-tubulin antibodies. (B) Genotyping two of AMBRA1 MDA-MB-231 cell lines. Deletions are underlined. Predicted amino acids are shown under the deleted nucleotide sequences. Extra introduced amino acids by the frame shift are highlighted in red. (C) qPCR analysis of AMBRA1 mRNA levels in control and AMBRA1-KO MDA-MB-231 cells.

**Figure S2.**
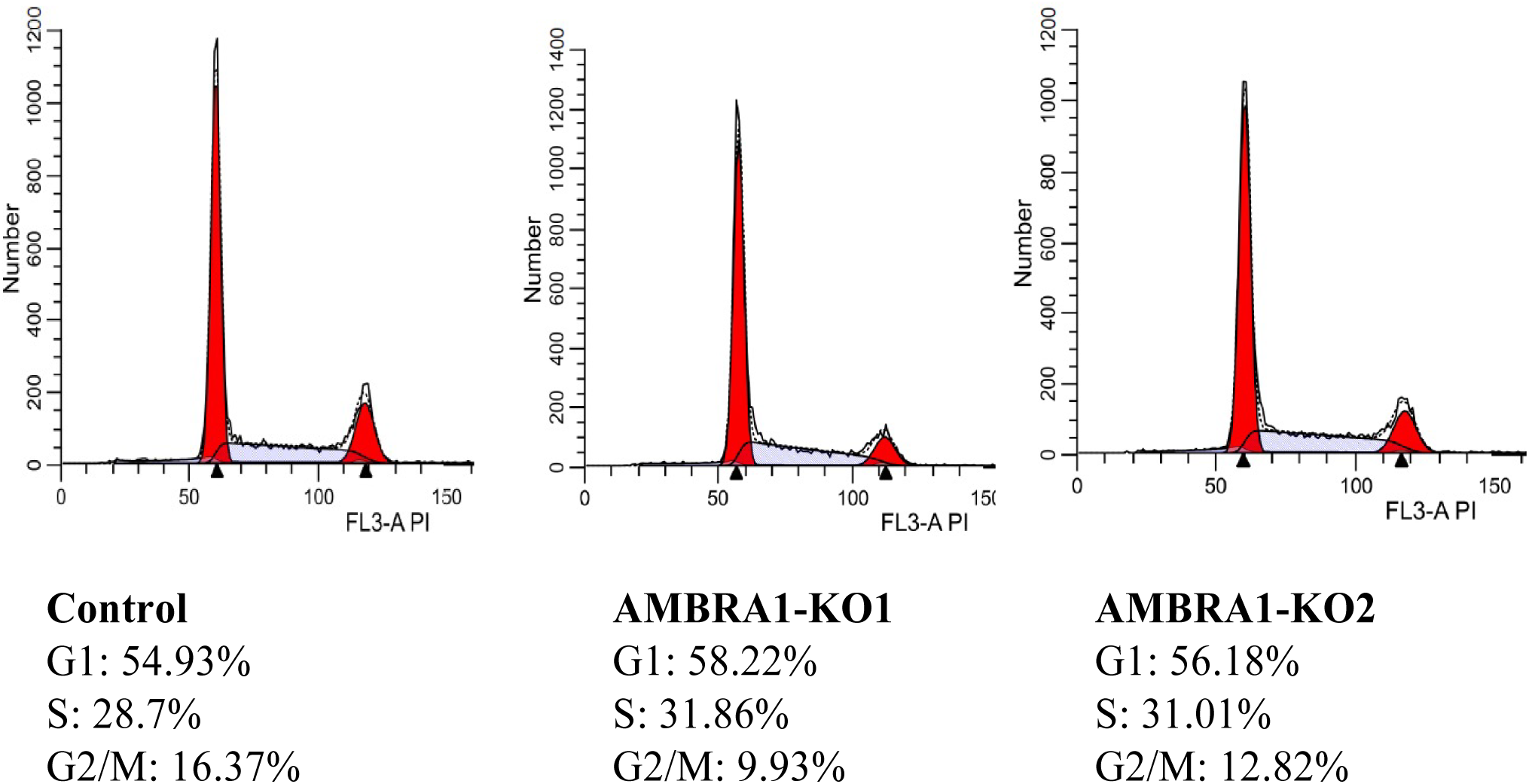
Control and AMBRA1-KO MDA-MB-231 cells were analyzed for DNA content by flow cytometry. Cells were fixed, permeabilized, and stained with propidium iodide (PI) and the cell cycle distribution of each cell type was performed by flow cytometry (see Materials and methods). Quantitative cell cycle phases proportions were identified by calculating the cell number % of each cell cycle phase relative to total phases after appropriate gating of cell populations by PI fluorescence and showed under the histogram of DNA content distribution.

**Figure S3.**
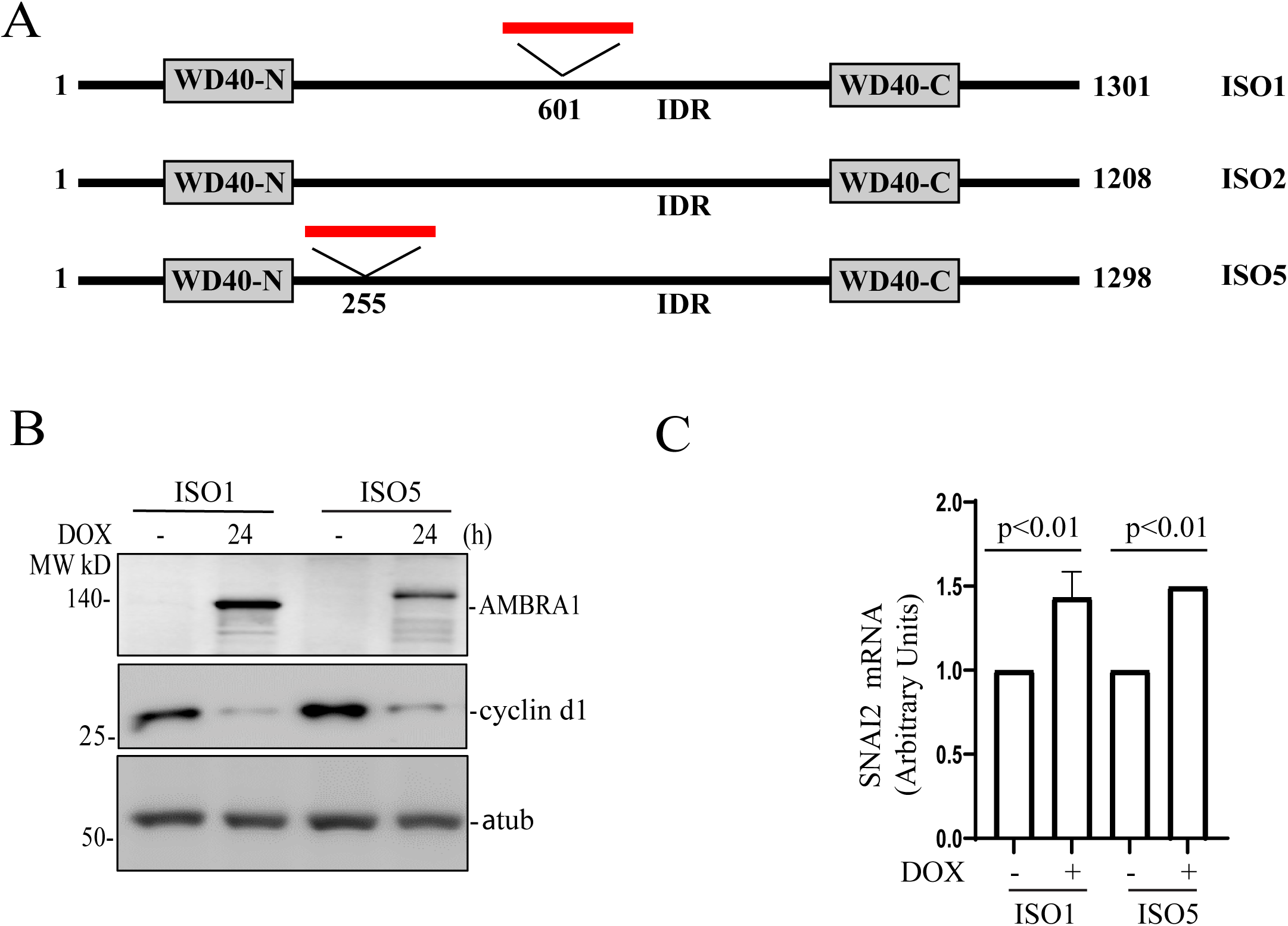
(A) A schematic representation of AMBRA1 isoforms 1, 2, and 5. The short lines in red represent insertions. (B) AMBRA1-KO MDA-MB-231 cells stably expressing dox-inducible AMBRA1 isoforms 1 or 5 were incubated with 100 ng/ml doxycycline for 48h. Cell lysates were immunoblotted with anti-AMBRA1, anti-cyclin d1, or anti-α-tubulin antibodies. (C) qRT-PCR analysis of the levels of SNAI2 mRNA in AMBRA1-KO MDA-MB-231 cells stably expressing either dox-inducible AMBRA1 isoform 1 or 5. The means and SD were calculated from three independent experiments.

**Figure S4.**
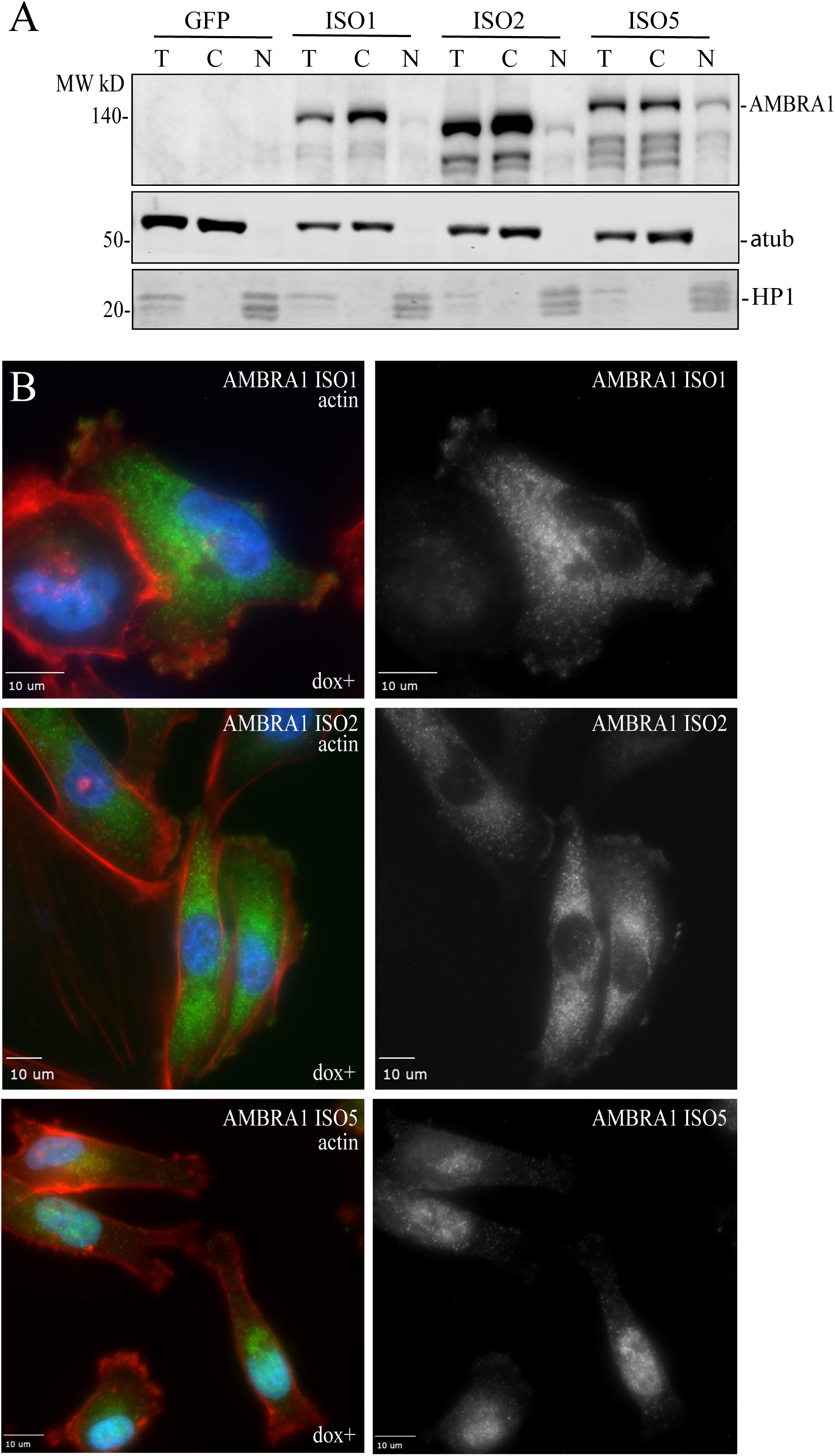
(A) AMBRA1-KO MDA-MB-231 cells stably expressing dox-inducible GFP, AMBRA1 isoforms 1, 2, or 5 were incubated with 100 ng/ml doxycycline for 48h. Total (T), cytoplasmic (C), and nuclear (N) fractions were collected (see Materials and methods) and subjected to Western blot by indicated antibodies. (B) AMBRA1-KO MDA-MB-231 cells stably expressing dox-inducible AMBRA1 isoforms 1, 2, or 5 were plated on collagen-coated coverslips for 24 hours and then were incubated with 100 ng/ml doxycycline for 24h. Cells were then fixed and stained with phalloidin-Alexa Fluor 594 (red) and anti-AMBRA1 antibodies (green).

